# A survey of influenza subtypes in olive baboons in selected areas in Kenya

**DOI:** 10.1101/380345

**Authors:** Emmanuel Kulwa Bunuma, Lucy Ochola, Andrew K Nyerere

## Abstract

3.

**Background:** Worldwide infections with influenza A viruses are associated with substantial illness and death among mammals and birds, in humans it accounts for 250,000-500,000 deaths per year its continuous mutation in different hosts poses a threat that can result in the emergence of a novel virus with an ability to cause a widespread pandemic. Surveillance of Influenza A viral genome from diverse hosts and subtyping is critical in understanding of the antigenic shift and drift of the influenza virus especially in hosts that are closely related to human beings like the Non-Human Primates (NHPs),pigs and birds. This study therefore identified the influenza subtypes circulating in *Papio anubis* (Olive baboons) at the interface of human and NHPs in Kenya.

**Methods:** Fifty nasal swabs samples were collected from baboons from the colony at the Institute of Primate Research (IPR), these animals were originally collected from Olorbototo, Yatta, Aberdares, Movoloni and Laikipia. The nasal swabs were collected in viral transport media using sterile dacron swabs and stored at −80°C. In this study, samples were screened initially using real time RT-PCR-CDC protocol for influenza A virus detection that targets the matrix gene and twenty five were found to be positive.

**Results:** The proportion positive were as follows, Olorbototo (75%), Ngurumani (44%) Aberdares (43%), Mavoloni (37.5%), Yatta (14%), and Laikipia (9%). These samples were taken through conventional PCR to amplify the haemagglutinin, neuraminidase and the matrix genes and eight samples were successfully amplified and later sequenced through 24-capillaries ABI 3500 XL Genetic Analyzer.Upon BLAST of these sequences, influenza subtypes H1N1 and H3N2 were detected. It was observed that the subtypes in baboons were as follows Olorbototo H1N1,Yatta H3N2, Aberdares H3N2, Mavoloni H1N1, Ngurumani H1N1 and Laikipia H1N1.Upon further analysis, the influenza positive Olive baboons were found to have been reared in the colony at at IPR colony for between 1-2 years and were in close contact with personnel.

**Coclusion:** Given the presence of H1N1 and H3N2 subtypes in baboons suggests that baboons can be naturally infected with seasonal endemic human influenza viruses, avian emerging pandemic or pandemic swine flu origin.

## 5. Introduction

Globally, influenza A virus infection is accompanied with significant sickness and death amongst mammals and birds (Karlsson et al. 2012), in humans it is responsible for up to 5 million severe cases and between 250 000 to 500 000 deaths worldwide per year (Who et al. 2009).The health of humans and animals is largely interlocked with 6 out of 10 emerging diseases originating from animals (Reperant et al. 2016). Many dynamics in relation to animals, environments and humans lead to the emergence of zoonotic diseases. The environments linked with pathogens and their reservoir hosts are continuously mutable and the rate of change is growing. The drivers of change include the transformation of farming practices, predominantly in the unindustrialized world, habitat obliteration; human invasion and climate change (Morens & Fauci 2013).

It is critical to assess and understand the influences of these changes on the interfaces between Influenza type A and their hosts, and amongst the host and other species, as well as other wildlife, livestock and humans (Fuller et al. 2013). Nonhuman primates, the closest living relatives of humans, are susceptible to other respiratory viruses like paramyxoviruses that cause respiratory disease in humans (Sasaki et al. 2013). These considerations prompted additional searches for species harboring novel influenza in baboons.

Mammals such as baboons are likely conduits for cross-species transmission of respiratory pathogens like influenza viruses because of their close and long-term contact with their owners, scientists, audiences, domestic animals, wild animals, and birds. (Fuller et al. 2013) Identifying the Influenza subtypes circulating in baboons at the interface of human to non-human primates was the aim of this study with the null hypothesis that the Kenyan baboons at the human-animal cannot be infected with Influenza virus.

## 6.0 Methods

### 6.1 Study site

The study was conducted on purposively wildly caught baboons from Olorbototo (Ngurumani), Aberdares, Ngurumani, Omolon (Oldonyo sabuki), these baboons had stayed for 1-2 years at the Institute of Primate Research at the time of this study. Figure 1 shows the location of the sites where the animals were caught within Kenya.

**Figure 1.**
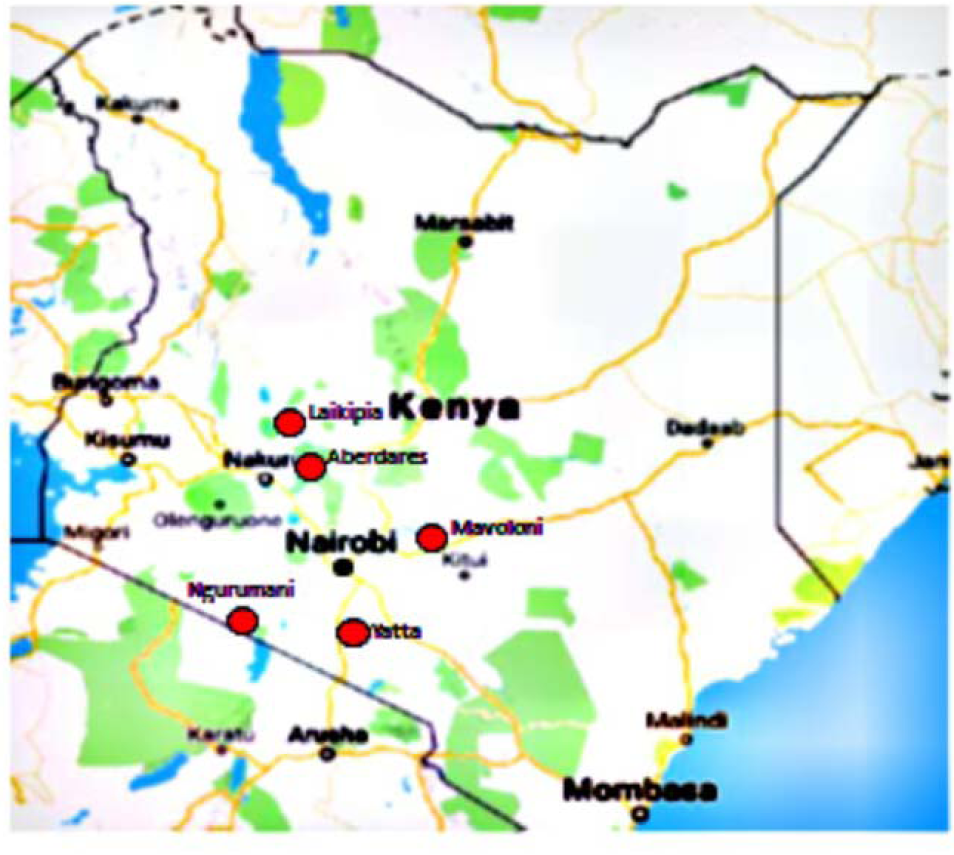
Map of Kenya showing the sites where the animals originated from.

### 6.2 Study design

This was a cross-sectional study carried out on fifty nasal swab samples collected purposively from wild caught Olive baboons (*Papio anubis*) kept in the colony at the Institute of Primate Research (IPR) (1°20′47.95″S, 36°42′51.13″E, Nairobi Kenya), where the presence and subtypes of influenza virus circulating in baboons from selected sites in Kenya was determined.

### 6.3 Ethics statement

All procedures reported herein were performed in accordance with institutionally approved animal care and use protocols with reference number IERC/08/16 approved by the Institutional Scientific and Ethics review Committee of the Institute of Primate Research Kenya. The committee is guided by the institutional guidelines as well as the international regulations including those of WHO and Helsinki conventional on the humane treatment of animals for scientific purposes and GLP. (Appendix 1)

### 6.4 Clinical assessment of animals

Animals within the colony at IPR undergo physical examination daily. Information relating to physical and bio data collected are indicated in Appendices 3,4 and 5. These clinical assessments were compared to the available classical flu fever signs in human for host response variability between non-human primates to the response seen in humans.

### 6.5 Laboratory methods

#### 6.5.1 RNA extraction from the viral nasal swabs samples

The virus RNA was extracted from the clinical nasal swabs of Baboons using the QIAamp^®^ Viral RNA extraction kit (Qiagen, Germany) following the manufacturer’s protocol. Briefly, 150μl of the viral transport media was added to 500μl of lysis buffer per tube and allowed to incubate at room temperature for 10 min to allow for the lysis.

A 500μl aliquot of ethanol was added and pulse vortex performed for 15s to give a homogeneous solution. A 650μl volume of the lysed solution was put to the spin columns and centrifuged at 8000 G for 1 minute and column placed in a clean collection tube. Then 500μl of Buffer AW1 was added to the spin column and centrifuged at 12000G for 1min in a Eppendorf 5415R centrifuge (Eppendorf AG, Barkhausenweg, Hamburg, Germany) and the column placed in a clean collection tube. Then the column was washed with 500μl of Buffer AW2 and centrifuged at 13,000 G for 3 min in a centrifuge.

Then finally spin column was positioned in a 1.5 ml micro centrifuge tube and 60μl of Buffer AVE added to the column and allowed to incubate at room temperature for 1 min. The column was then centrifuged at 8,000 G in the Eppendorf centrifuge at room temperature for 1 minute to the filtrate (RNA) then storage was done at −80°C.

#### 6.5.2 Real time RT-PCR

Real-time PCR amplification and screening was performed on an ABI 7500 (Applied Biosystems, CA, USA) using the AgPath-ID one-step RT-PCR kit (Applied Biosystems, Foster City, CA, USA). The total 25μl reaction volume for each sample enclosed 5 μl of extracted RNA, 12.5 μl of AgPath Kit 2X buffer, 1 μl of AgPath 25X enzyme mix, 5 pmol of Taqman probe, 10 pmol of each of the forward and reverse primers, and 6 μl of RNase-free water. Each RNA sample was tested by sets of matrix gene (Conserved gene across the subtypes) primers. For sample screening reverse transcription was achieved at 50°C for 30 min and 95°C for 15 min. PCR was achieved after 45 cycles of denaturation at 94°C for 15s and annealing at 55°C for 30s and final extension at 68°C for 5minutes the cycle threshold ≤40 was interpreted as positive.

#### 6.5.3 Conventional RT-PCR for genomic amplification

The RT-PCR was performed using Superscript II One-Step RT-PCR system Platinum ™ /Taq mix (Invitrogen Corporation, NY, USA). The reaction mix was organized by mixing 12.5μl of the 2x reaction mix, 0.5μl of the forward primer (20μM), 0.5μl the reverse primer (20μM) primers details and source are shown on the appendix 2-, 1.0μl Superscript II RT/Platinum Taq mix and this mixture was then capped using 7.5μl of distilled water to make a total of 22μl. 7μl of the RNA template was then added making the individual reaction volume to 29μl. Thermocycling conditions ware, 1 cycle of reverse transcription at 50°C for 30 min followed by an initial denaturation of 94°C for 2min. This was followed by 35 cycles of; denaturation at 94°C for 30s, annealing at 55°C for 30s and strand extension at 68°C for 1 min. Finally the reaction mixture was incubated at 68C for 1min to allow for extension of recessed ends of the amplicons. A final pause was set at 70 °C.

#### 6.5.4 Visualization of the amplicons by Agarose Gel electrophoresis

The agarose prepared was 1% and was prepared in 1x TBE buffer. The solution was mixed by spinning gently and then heating in a microwave until all the agarose melted.

Cooling was done for a few min at room temperature and then the gel was added at the ratio of 400mls of gel to 5μl of Ethidium bromide. Then the gel liquid at 37°c was then poured into an electrophoretic tank with a comb and left to set and solidify for 40 minutes at room temperature. The combs were then carefully removed. 5μl of the PCR samples were mixed with the 3μl of the blue orange gel loading dye (Invitrogen, NY, USA) and then loaded onto the wells. A 1kb DNA ladder marker (Invitrogen, NY, USA 10787-018) was loaded on the first lane of each of the wells. The tank was run at 60 volts for about 90 minutes. The gel was then visualized and observed on the Alpha Imager gel documentation system (Alpha Innotech, CA, USA).

#### 6.5.5 Clean-up of PCR products using Exosap-IT

Removing the dNTPs and primers from PCR product was done. PCR tubes covering 10μl of the PCR products to be purified were shortly spun then 3 μl of the ExoSap-IT enzyme (U. S Biological, Swampscott, MA, USA) was added to each of the PCR tubes, then followed by a brief vortex for 10s. They were then spun for 30s. The PCR tubes were then placed into Thermal Cycler, and incubated for 30 min at 37°C. Then inactivation of the ExoSap-IT enzyme by incubation for fifteen minutes at 80 °C before storing the product at 4 °C.

#### 6.5.6 Cycle sequencing of the purified PCR products

PCR amplicons including fluorescent-labeled dideoxy-chain terminators were created using an ABI BigDye Terminator version 3.1 cycle sequencing Kit (Applied Biosystems, Forster City, USA). The reaction mixture for both the forward and reverse reactions were organized by adding 2μl of BigDye to 2μl BigDye 5X buffer then finally by addition of 1μl (4μM) of the M13R/F primers and 3μl of distilled water to make a total volume of 8μl. This reaction mixture was then loaded into each annotated well on the 96-well plate followed by addition of 2μl of the purified PCR product. The plates were enclosed with a sealing mat and were vortexed briefly.

The PCR running conditions were 1 cycle of initial denaturation at 95 °C for 5 min, followed by 30 Cycles of denaturation at 95 °C for 15 s, annealing at 45 °C for 30 s and strand extension at 68 °C for 2 min and 30 s. This was followed by a final incubation at 68 °C for 3 min to allow for extension of settled ends of the amplicons before storing the product at 4 °C.

#### 6.5.7 Purification of the cycle sequencing products using sephadex spin columns

®Sigma Dry sephadex G-50 medium powder was loaded into unused clean wells of 96-well Column Loader (Millipore, MA, USA). The 96 well Multi-Screen®-HV Plate Millipore was then placed upside down on top of the column loader, and both the Multi-Screen Plate and the column loader were held together and upturned. The top of the Column loader was selected to release the resin. Then 300μl of Milli-Q water was then added to each well encompassing sephadex to swell the resin and the setup allowed to incubate at room temperature for 3 hours. The Multiscreen HV plate was then placed on top of a standard 96-well micro plate and then was spun at 8000rpm for 5 min on an Eppendorf 5810R bench top centrifuge (Eppendorf, Hamburg, Germany) using a two-tray rotor to eliminate extra water from the columns.

Extra water was then castoff. The sequencing products were sensibly added to the centre of each Sephadex well. The 96-well plate with the excess water was replaced with a new 96-well microplate (USA Scientific, FL, USA), and centrifugation done at 910× g. for 5 min guaranteeing that approximately 10μl of product came through the column. Then, 10μl of Hi-Di™ Formamide (Forster City, CA, USA) was added to guarantee the sequencing fragments were upheld as single strands and to hydrate any dry (empty) wells to circumvent extinguishing the capillaries.

#### 6.5.8 Genetic analyzer procedure

The filtered PCR products from samples PAN 3837,PAN3342,PAN3350,PAN 3340,PAN 3638,PAN 3630, PAN 3201 and the Hi-Di in the 96 well plate were placed into 24-capillaries ABI 3500 XL Genetic Analyzer (Applied Biosystems). These were left to run and nucleotide sequences were obtained using the sequence analysis software (Applied Biosystems).

#### 6.5.9 Contiguous assembly

To make contiguous nucleotide sequences from the reverse and forward sequence runs for each amplified segment from the Genetic Analyzer, the sequences were put into the contig assembly program (CAP) of DNA Baser Sequence Assembler v3 (Heracle BioSoft SRL Romania, http://www.DnaBaser) and the consensus sequences were generated for BALST at https://blast.ncbi.nlm.nih.gov.The sequences are shown at the appendix =

#### 6.5.10 Similarity searches

To determine whether the obtained nucleotide sequences were similar to Influenza A sequences deposited in genomic databases Blastn was performed at https://blast.ncbi.nlm.nih.gov.

### 6.6 Biosafety measures taken

Biosafety Level (BSL) 3 cabinet was used with protective equipment’s worn all the time when handling the samples which includes laboratory coat, gloves and face masks, appropriate sample labelling, washing hands before and after every laboratory work.

### 6.7 Limitations of the study

This study could not culture the unsubtyped baboons influenza virus in avoidance of the risks that might have been associate with it, and hence this made very low nucleic acid materials available for different target gene amplifications this factor also limited this study from doing serological tests as preliminary test and hence opted the real time PCR as the screening method as recommended by (WHO, 2012),limited time allocated for the study could not allow sampling of other species and hence data for avians and humans for this study are missing.

## 7.0 RESULTS

### 7.1 Real time Reverse Transcriptase PCR

Real time RT-PCR of the Matrix gene was used to screen fifty nasal swab samples collected from the colony. Of the fifty samples we found that 25 were positive by RT-PCR (on the appendix 7) it was reported that the colony baboons had stayed for 1-2 years at IPR. Results further indicated that majority of the positive animals were from Olorbototo (75%), Ngurumani (44%), Aberdares (43%), Mavoloni (37%), Yatta (14%) and Laikipia 9% (Table 1). This test was done in animals that ranged from 1–10 years of age at the time of sampling. Juvenile male baboons 1-4 years normally reach 5-7kgs in weight, while sub adult male 7-10 years reach 14-15kgs, juvenile female 3-4kgs and adult females reaches up to 13kgs. The results on the real time RT-PCR shows that juveniles are mostly infected compared to adults and sub-adults and that the total numbers of males infected were 11 and females were 12.

**Table 1:** The results for the real time RT-PCR

| **Sample ID** | **CT Value** | **Area of collection** | **Sex** | **Weight** | **Age** |
| --- | --- | --- | --- | --- | --- |
| PAN 3642 | 41.091 | Olorbototo | M | 16 | 6 |
| PAN3637 | 40.513 | Olorbototo | M | 18 | 7 |
| PAN 3837 | 38.1461 | Yatta | F | 12 | 5 |
| PAN 3342 | 38.605 | Olorbototo | F | 9 | 4 |
| PAN 3641 | 42.03 | Olorbototo | M | 19 | 6 |
| PAN 3625 | 44.105 | Olorbototo | F | 14 | 5 |
| PAN 3629 | 40.157 | Olorbototo | F | 15 | 6 |
| PAN3188 | 41.20 | Yatta | F | 14 | 5 |
| PAN 3340 | 38.099 | Olorbototo | M | 12 | 5 |
| PAN3638 | 34.219 | Olorbototo | M | 9 | 4 |
| PAN 5536 | 36 | Yatta | M | 16 | 6 |
| PAN3630 | 37.1206 | Aberdares | F | 15 | 5 |
| PAN 3201 | 38.9381 | Olorbototo | F | 10 | 5 |
| PAN3484 | 44.032 | Yatta | M | 9 | 6 |
| PAN 3585 | 42.502 | Olorbototo | M | 13 | 5 |
| PAN 3315 | 37.1621 | Mavoloni | M | 11 | 6 |
| PAN 3197 | 43.102 | Olorbototo | F | 10 | 4 |
| PAN 3587 | 41.323 | Olorbototo | F | 9 | 5 |
| PAN 3335 | 42.012 | Olorbototo | M | 19 | 6 |
| PAN 3350 | 38.4184 | Olorbototo | F | 21 | 6 |
| PAN 3632 | 40.371 | Olorbototo | F | 17 | 5 |
| PAN 3647 | 35.9276 | Olorbototo | F | 14 | 4 |
| PAN 3623 | 43.812 | Olorbototo | F | 11 | 6 |
| PAN 3624 | 37.2694 | Olorbototo | F | 12 | 5 |
| PAN 3464 | 44.353 | Olorbototo | F | 24 | 7 |
| PAN 3613 | 39.5136 | Aberdares | M | 18 | 6 |
| PAN 3459 | 40.323 | Olorbototo | M | 23 | 6 |
| PAN 3181 | 38.8304 | Aberdares | M | 15 | 4 |
| PAN 3540 | 37.157 | Yatta | M | 13 | 5 |
| PAN 3560 | 45.392 | Orbototo | F | 16 | 6 |
| PAN 3646 | 39.3754 | Orbototo | M | 17 | 5 |
| PAN 3479 | 48.2390 | Olorbototo | F | 10 | 4 |
| PAN 3334 | 37.7283 | Aberdares | F | 11 | 5 |
| PAN 3538 | 47.293 | Mavolani | F | 9 | 4 |
| PAN 3343 | 43.393 | Orbototo | F | 10 | 4 |
| PAN 3339 | 38.3343 | Mavolani | F | 11 | 5 |
| PAN 3186 | 38.5032 | Mavoloni | F | 12 | 5 |
| PAN 3631 | 44.282 | Mavoloni | M | 13 | 4 |
| PAN 3505 | 37.6242 | Mavoloni | M | 14 | 5 |
| PAN 3621 | 43.4492 | Yatta | M | 15 | 4 |
| PAN 3643 | 35.1316 | Olorbototo | M | 16 | 5 |
| PAN 3837 | 37.4909 | Laikipia | M | 18 | 6 |
| PAN3338 | 38.2143 | Yatta | M | 15 | 5 |
| PAN 3640 | 38.5427 | Ngurumani | F | 13 | 4 |
| PAN 3537 | 41.34334 | Ngurumani | M | 15 | 5 |
| PAN 3634 | 46.325 | Olorbototo | M | 16 | 6 |
| PAN 3201, | 38.5026 | Ngurumani | F | 18 | 7 |
| VE + | 28.17 |  |  |  |  |
| VE _ | _VE |  |  |  |  |

### 7.2 Amplification of the positive samples from real time rt-PCR

The RT-PCR one step for HA fragment amplification was done in PAN3837, PAN3342, PAN3350, PAN3315 nasal swabs samples whereas the NA fragment amplification was done in PAN 3340, PAN3338, PAN3640 and PAN3201, finally the NA fragment was successfully amplified in PAN3350 and PAN3315 and Matrix genes was successfully amplified in PAN3640 and PAN 3201. The reason for missing bands and lack of multiple amplification of samples were due to low volume of nucleic acid present and hence unable to culture unsubtyped novel influenza virus from the baboons, the bands obtained are displayed on gel photos (Figure 2 and 3).The image was observed on Alpha Imager.

**Figure 2:**
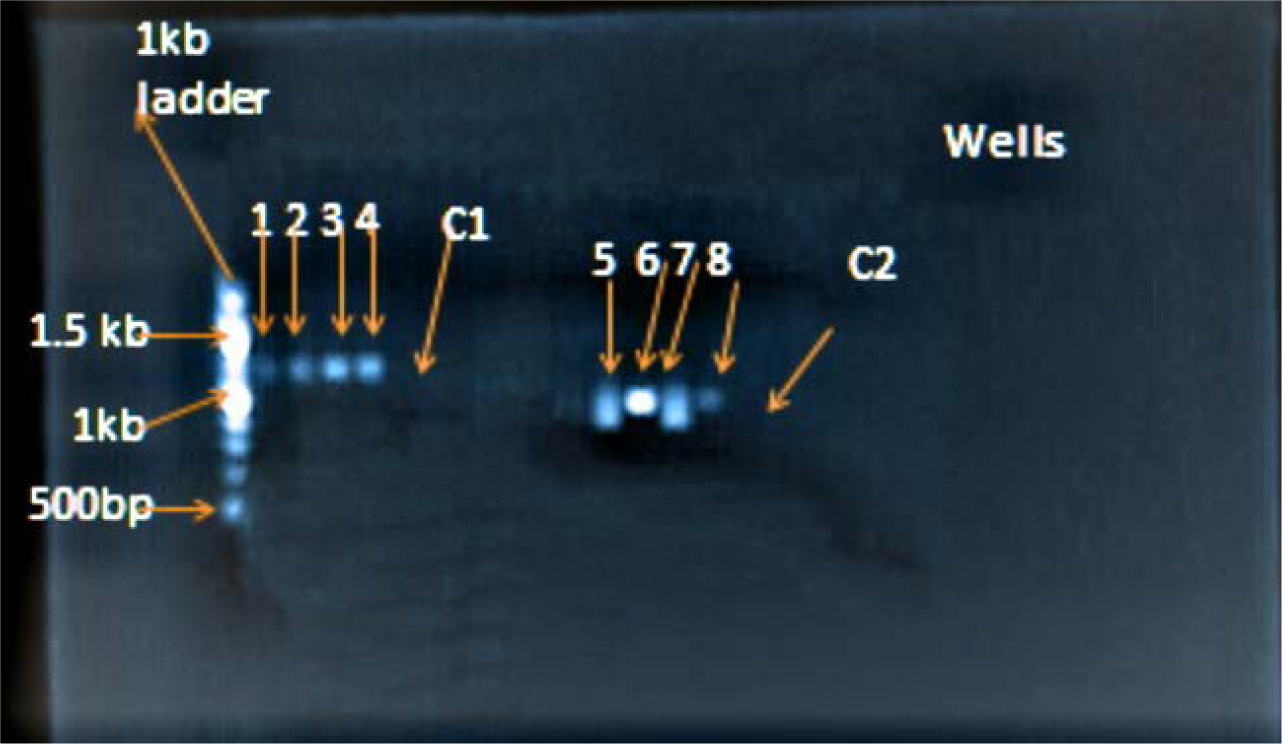
Gel photo showing the PCR amplification of the three influenza gene (HA, NA, genes) Segments. Lane 1, 1kb ladder marker, Lane 2, 3, 4, 5, is HA fragment for PAN3837, PAN3342, PAN3350, PAN3315 respectively, Lane 6 is negative control for HA, Lane 7,8,9,10, is NA fragments for PAN 3340, PAN3338, PAN3640 and PAN3201 respectively, Lane 11 is negative control for NA.

**Figure 3:**
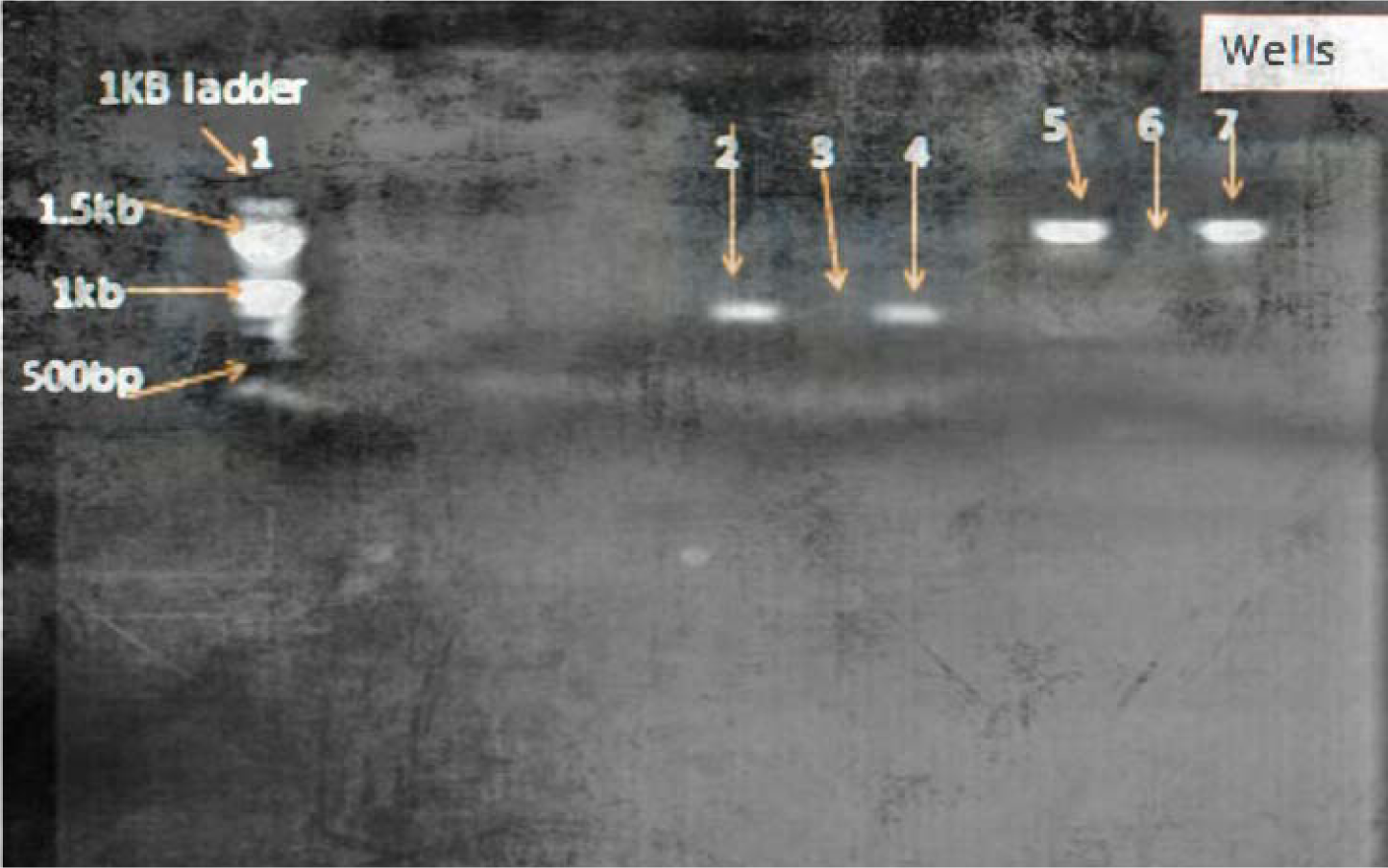
Gel photo showing the PCR amplification of the three influenza gene ( NA, and M genes) Lane 1, 1kb ladder marker, Lane 2 and 4 are NA fragments for PAN3315 and PAN 3350 respectively and Lane 3 is the Negative control, Lane 5 and 7 is MA fragment for PAN3640 and PAN 3201 respectively and lane 6 is the negative control.

### 7.3 Sequences similarity searching in the gene bank

To conclude whether the obtained nucleotide sequences from the Kenyan baboons were similar to other influenza sequences placed in genomic databases, a similarity search against sequences in the influenza virus resource was done using blastn (http://blast.ncbi.nlm.nih.gov/) (Appendix 8) which compares the query to the other sequences deposited in the Influenza genome database.

### 7.4 Clinical assessment of influenza positive *P. anubis*

The age of the animals at the time of nasal swab collection was estimated to be between 1-10 years. Of these 80% of the animals were between 1-5 years of age and these were also positive for influenza infection Using amplification for the matrix gene by real time RT –PCR it was found that the positive samples were from Olorbototo (75%), Ngurumani (44%) Aberdares (43%), Mavoloni (37%), Yatta (14%), and Laikipia 9%.

These samples were taken through one step RT-PCR and only eight samples gave bands, which were then run through the 24-capillaries ABI 3500 XL Genetic Analyzer for sequencing and gave the following subtypes in baboons from Olorbototo H1N1,Yatta H3N2, Aberdares H3N2, Mavoloni H1N1, Ngurumani H1N1 and Laikipia H1N1.

The clinical signs of these subtyped baboon influenza isolates were documented as per the clinical information forms attached on appendices 5, 6 and 7, and presented on Table 4.2 below.

**Table 2:**
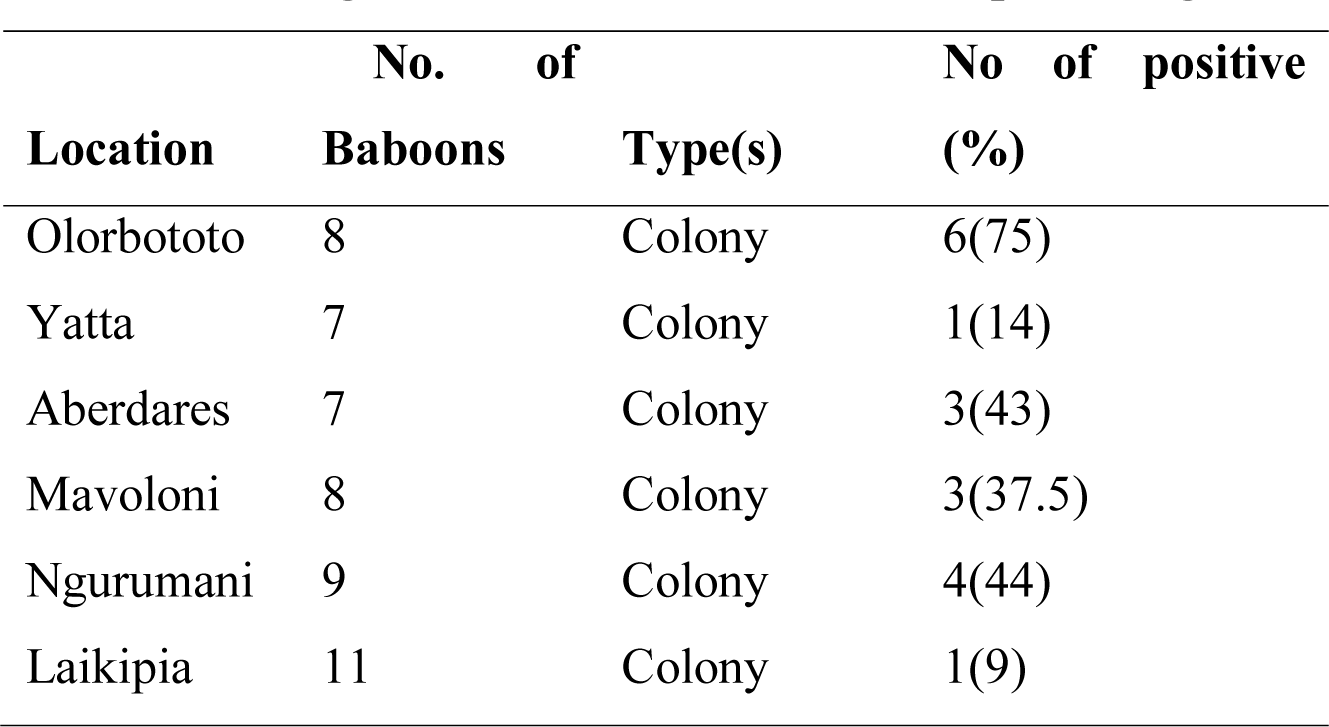
The original location of *P. anubis* and percentage of influenza positive samples.

**Table 3:**
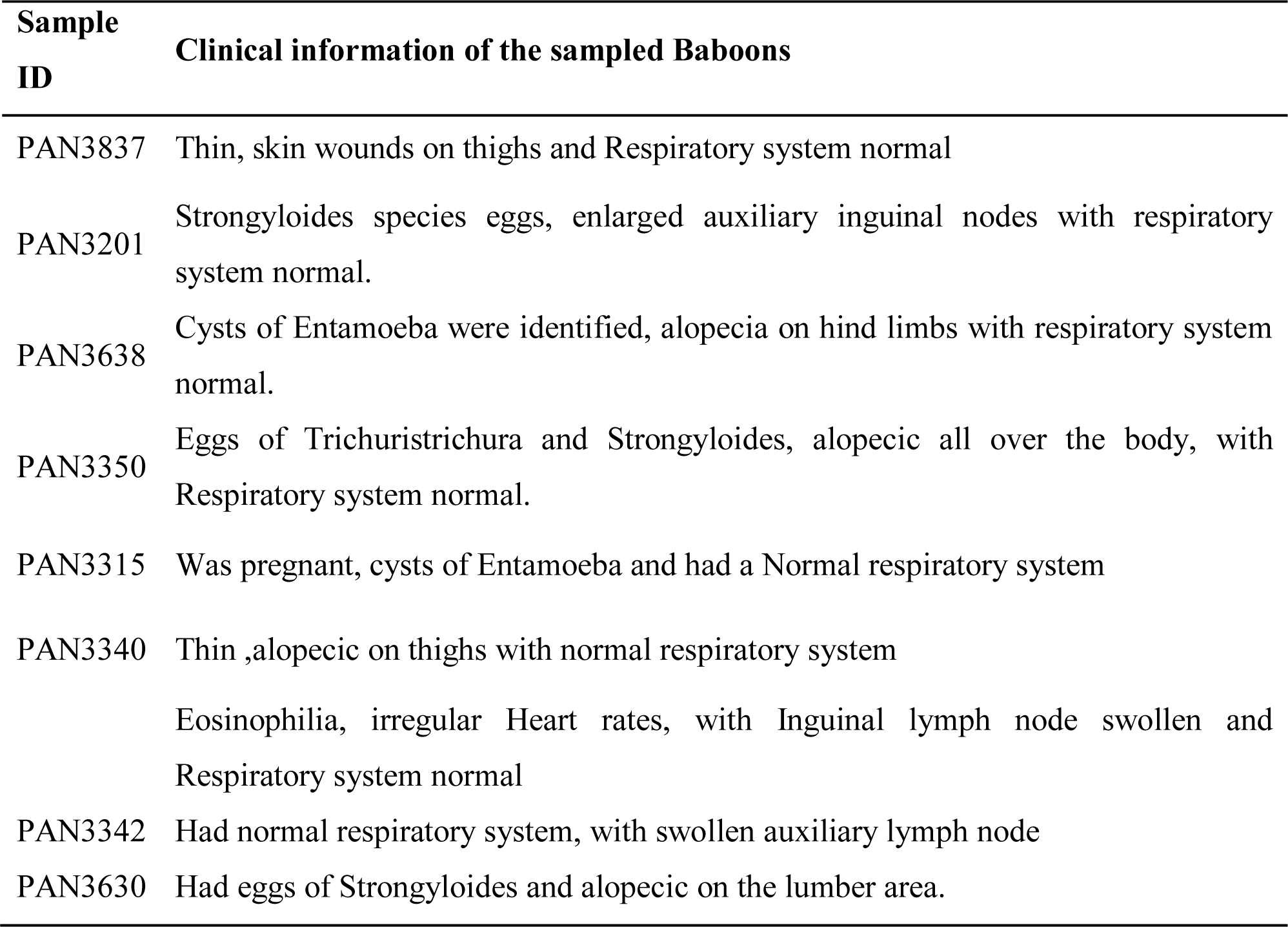
Clinical symptoms of baboons found to be positive with influenza A but yet they didn’t present the clinical influenza fever symptoms

## 8.0 Discussion

In Africa Non-Human Primates are widely distributed for example in Zambia Baboons and vervet monkeys live side by side with humans in game management areas and this situation often leads to high levels of human–baboon/monkey conflicts, in these settings, scientists have found Human Parainfluenza type 3 from Humans in baboons,(Sasaki et al. 2013). Human metapneumo virus in wild great apes in Rwanda (Sasaki et al. 2013). In Kenya, studies have shown the presence of influenza in dogs, but there is no data available on baboons yet there are found in large numbers and are widespread. Based on this, the baboons from a selected area in Kenya were screened to identify the influenza subtypes circulating in them.

Upon screening of fifty samples of the wildly caught colony kept baboons, 25 were found to be positive for the M-gene, of which 80% of the positive samples were from juvenile animals (1-5 years).These results are consisted with (Karlsson *et al.,*2012) who screened 48 non-human primates and found 14 positive animals, In 2005, Whittier *et al*. reported positive titers to influenza A and B viruses as part of a wider survey of seroprevalence of infectious agents in non-human primates. This shows that non-human primates are also infected by influenza virus. Despite baboons being infected with influenza it was observed that they lack clinical signs, this was in agreement with earlier studies that observed the same that even though the viruses replicate well in the respiratory tract, animals do not generally develop any symptoms of disease (Margine *et al.*, 2014) contrary of what is seen in humans, normally accompanied with pneumonia acute respiratory failure and runny nose (Leslie et al., 2016) which is frequently complicated by bacterial co-infection This may suggest that baboons are either opportunistic hosts or infections with influenza viruses in baboons might exacerbate the development of other diseases such as active TB which they succumb to (Tarara *et al,* 1985). Viral infections induces the type I interferon’s which inhibits the interferon-γ mediated immune responses which works to inhibit the development of active Mycobacterium tuberculosis (De Paus *et al,.* 2013).

Two subtypes H1N1and H3N2 were identified in this study after sequencing and blastn. The H5N1 and other subtypes from avian species were not found which might indicate there were no avian species contact flu transmission. These observations further confirm results from an earlier study where seasonal subtype H1N1 and H3N2 influenza A strains were detected in performing macaques at frequencies of in Cambodia (29.2%), Singapore (16.7%), Sulawesi (16.1%), Bangladesh (13.3%), and Java (6.0%). (Karlson *et al.,* 2013). This study observed the following subtypes in baboons from Olorbototo H1N1,Yatta H3N2, Aberdares H3N2, Mavoloni H1N1, Ngurumani H1N1 and Laikipia H1N1.

The presence of influenza infections H1H1 and H3N2 subtypes in non-humans primates in Kenya confirms that baboons can also be infected with Influenza virus nevertheless with non-classical flu fever. Extra epidemiologic studies of humans and wild caught colony kept nonhuman primates are needed to determine whether Influenza virus is transmitted between humans and wild nonhuman primates or from other species like the porcine and avian.

### 8.1 Conclusion

Screening of nasal swabs by real-time reverse transcription PCR results indicated the presence of the conserved matrix gene which confirms the presence of influenza viruses in the baboons. Non-human primates can therefore naturally be infected with influenza viruses and the viruses replicate in the respiratory tract but without necessarily developing the clinical signs. This means that they can harbor viruses unnoticed if we only depend on clinical signs to diagnose. The subtype H1N1 and H3N2 influenza A strains were identified circulating in non-human primates. These two subtypes have also been identified in human, so further epidemiologic studies can be done to search how baboons affect the transmission of these viruses.

### 8.2 Recommendations

There is a need for continuous surveillance and monitoring of genetic changes and virus evolution in wild and domesticated animal populations for example bats, rodents, camels. Continuous surveillance would also be critical in identifying other influenza viruses circulating in these animal populations in Kenya. Assimilating this surveillance with on-going surveillance for influenza viruses in humans will provide useful information for public health action vaccines, novel flu virus.

## 8.3 Acknowledgements

This work was supported by the Africa Union – African innovation JICA – JKUAT and PAUSTI joint Project.

## Acknowledgements

My sincerest gratitude to Prof. Wallace D. Bulimo from University of Nairobi who assisted in molecular analysis of this work.

## Partial sequences of the positive influenza nasal swabs isolates

### >1 H1N1 2016 Isolates from Baboons PAN 3837

TTATTTTTTTTTTTGTCCTCAGGGAGCAAAAGCAGGGGTCAGGATATGCAGCCGATCT

GAAGAGCACACAAAATGCCATCGATAAGATTACTAACAAAGTAAATTCTGTTATTGA

AAAGATGAATACACAGTTCACAGCAGTGGGTAAAGAGTTCAACCACCTTGAAAAAAG

AATAGAGAATCTAAATAAAAAAGTTGATGATGGTTTCCTGGACATTTGGACTTACAAT

GCCGAACTGTTGGTTCTACTGGAAAATGAAAGAAATTTGGACTATCACGATTCAAATG

TGAAGAACTTGTATGAAAAAGTAAGAAACCAGTTAAAAAACAATGCCAAGGAAATTG

GAAACGGCTGCTTTGAATTTTACCACAAATGCGATAACACATGCATGGAAAGTGTCAA

GAATGGGACTTATGACTACCCAAAATACTCAGAGGAAGCAAAATTAAACAGAGAAAA

AATAGATGGAGTAAAGCTGGAATCAACAAGGATCTACCATATTTTTCGTCTCAGGGAG

CAAAAGCAGGGGGACATAGTGTTGGTAATGAAACGAAAACGGGACTCTAGCA

TACTTACTGACAGCCAGACAGCGACCAAAAGAATTTGGGATGGCCATCAAATAGTGT

ACGTTACTTTGTA

Similar to Influenza A virus (A/Shimoga/MCVRAG4823/2017(H1N1)(H1N1)) segment 4 hemagglutinin (HA) gene, partial cds.

### >2 H1N1 2016 Isolates from Baboons PAN 3350

ATAATAAAGTCTCTGTGTTGAGGAATGTGCATCCTCAACATCCTCGGGCT

CCTGCTTTTGCTCCCGGAGACGAATAAACCACAAAAGGATATGCTGCTCC

CGCTAGTCCAGATTGTGTTCTCTTCGGGTCGTCCTCTTATTAGTTCAACC

CAGAAGCAAGGTCTTATACAATCCAGCCCTGTTAGTTCTGGATGCTGAAC

AAAACTCCTGCTTTTGCTCCCTGAGACCAATAATACTACGATATCTTGCT

TTATTGAGAATTTATTGTCAGTCCCAGTCCATCCATTCGGATCCCAAATC

ATCTCAAAACCTTTTCTTGAACTAATGCTCTTAGTTCTCCCTATCCAAAC

ACCATTGCCGTATTTGAATGAAAATCCTTTTACTCCTGCTTTTGCTCCCT

GAAACCAATATAAA

Similar to Influenza A virus (A/Pennsylvania/56/2018(H1N1)) segment 6 neuraminidase (NA) gene, complete cds

### >3 H3N2 2016 Isolates from Baboons PAN 3340

TTCCAAAAAGCGCAGGAGCATATATAAGCGTGACATTGGCGTCCCCCCCA

TCGGGCCGTGCCTCTGTTCCTTCTGTCCCTGAAGTGCCACAAACTCAAGA

TTTCTGTTTGAGGTCCACAAGTTTTCTTTTTCCTCTTTTCTTCCCCTAAT

CAACTCAACATAAAAGCACCGATTGATGCATTTTTTGCCTTCAACAGAAA

AAATACCAAAATACCCGGACCTATCAAGAGGGAAATTTGTGAGAAGAATC

ATATTGTGAAATTGGAGAGAGAAACAGACCCAATCGTTTTTATCTTTTGA

TTGGGATTCATCTTTACCCCTGTTTTTGCCCCCAGAGAAAAAAAAACACA

CGTCTTTTCCATCATCAAATGCCCACCCTTTCACTCCTTGAACCCCTTCT

TCATTGATAAAATAAAAACAATGACTACTGCTGGAGCTGTCTTTTTTTCG

GGGTGTGTCTCCAACAAGCCCTGAACACACATAACGGAAAACAATGCTAT

GACCCTTTATGTTTATATCTACTAGGGGCCGATTGGATCCTTTCCATTTG

TCTCTGCAAACACCTCTGACACCGGGATATCGGGGATAGAAAGAGCTCTC

TTCCACATGCTGAGCACTTCCTGACTATGTGCTAGTATAAAAAATTCTCC

CCTCCTCAGAGAATAATTTTATAGTATCAGCTTTTCCTGTGGCTTTTCTA

TCAGTCTTCTCTACTGTACAATTACTATTGATACAAACGCTTTCAGACTC

CTGGTTTCGTGATCATCTCTTGAAAACGATGAAAAAAACACTATCACAAA

CT

Similar to Influenza A virus (A/Kenya/001/2017(H3N2)) segment 6 neuraminidase (NA) gene, complete cds

### >4 H1N1 2016 Isolates from Baboons PAN 3315

AAAAGTTCGACACTAATTGATGGCCATCCGAATTCTTTTGGTCGCTGTCT

GGCTGTCAGTAAGTATGCTACAGTCCCGTTTTCGTTTCATTACCAACACT

ATGTCCCCCTGCTTTTGCTCCCTGAGACGAAAAATCTGGTAGATCCTTGT

TGATTCCAGCTTTACTCCATCTATTTTTTCTCTGTTTAATTTTGCTTCCT

CTGAGTATTTTGGGTAGTCATAAGTCCCATTCTTGACACTTTCCATGCAT

GTGTTATCGCATTTGTGGTAAAATTCAAAGCAGCCGTTTCCAATTTCCTT

GGCATTGTTTTTTAACTGGTTTCTTACTTTTTCATACAAGTTCTTCACAT

TTGAATCGTGATAGTCCAAATTTCTTTCATTTTCCAGTAGAACCAACAGT

TCGGCATTGTAAGTCCAAATGTCCAGGAAACCATCATCAACTTTTTTATT

TAGATTCTCTATTCTTTTTTCAAGGTGGTTGAACTCTTTACCCACTGCTG

TGAACTGTGTATTCATCTTTTCAATAACAGAATTTACTTTGTTAGTAATC

TTATCGATGGCATTTTGTGTGCTCTTCAGATCGGCTGCATATCCTGACCC

CTGCTTTTGCTCCCTGAGAACGAAAAAAAAAAAAA

Similar to Influenza A virus (A/Shimoga/MCVRAG4823/2017(H1N1)(H1N1)) segment 4 hemagglutinin (HA) gene, partial cds

### >5 H3N2 2016 Isolates from Baboons PAN 3630

CTCAAGTTGCGAAGGCTTATATAAGCCTGACATTGACGTCCGCCCCATCA

GGCCATGACCCTGTTCCATCTGTACCTGAAGTGCCACGAACCACAAGATT

ACGGTTTGAGGTCCACAAGACTTCAGTTTCCTCTTTTCTTCCCCTAATCA

ACTCAACATAAAAGCACCGATTGATGCAGCTTTTGCCTTCAACAGAGAAA

ATACCAGAATAACCGGACCTATCAAGATGGCAATTTGCATGAAGAAGCAT

ATTGTGGAAATGGTGAGAGAAACAGAGCCAATCGTTATTATCTTTTGATT

TGGATTCATCTTTACTCCTGCTTTTGCTCCCTGAGACCAATAACCACACG

TCATTTCCATCATCAAATGCCCAGCCTTTCACTCCATGAACACCTTCTTC

ATTGTTAGAATTCAAACAATGACTACTGCTGGAGCTGTCGTTTTTTCTGG

GTGTGTCTCCAACAAGTCCTGAACACACATAACTGGAAACAATGCTATGA

TCCTTTATGTTTATATCTACTATGGGCCGATTGGATCCTTTCCAGTTGTC

TCTGCAGACACATCTGACACCAGGATATCGAGGATAGCAAGAGCACTCTT

CGACATGCTGAGCACTTCCTGACAATTTGCTAGTATGAACGATTTTCCCC

TCCTCAATGAATAGTATTTTAGTATCAGCTTTTCCTGTAGCATTTCCATC

AGTCATTACTACTGTACAAGTTCCATTGATACAAACGCATTCTGACTCCT

GGGTCCTGAGATATCTTTTGGAACATGAAAAACACTATCTACAAGCCTCC

CATTGTAAATGAAGCTAGCAGTTGCATTTTTATCATCCCCTGTTATACAA

ACATGCAGCCATGCTTTTCATCGTGACAACTTGAGCTAGGACCAAGGCTA

TTGCAACTGCCTGATCCAGAAATGGGAAGGAAACGCACCGAAGCACGACG

CTGCTGCTTGACGTA

Similar to Influenza A virus (A/Victoria/146/2016(H3N2)) neuraminidase (NA) gene, complete cds

### >6 H3N2 2016 Isolates from Baboons PAN 3342

CTTCTCCTTACTGGCATCCTTCCACAGCCAGAAGATTCGAGCAGAGAACTTGATGAGTACTTGGCCTGGTATAACGACAGATCTTGAAGACTCTCGATGGAACTGACATGCAAGACAAGATCAACTCTAGTCATCCTTCGACTTACGGGGATTTTATGGCCTGTTTTCACGGCTCACCGTGCCCAGTGAGCGAGGAGTGCAGCGTAGACTCCTTCGTCCAAAATGCCCTCAATGGGAATGGAGACCCAAATAACATGGACAAAGCAGTTCAAACTGTATAGGAAACTTAAGAGGGAGATAACGTTCCACGGGGCCAAAGAAATAGCTCTTAGTTATTCTGCTGGTGCACTTGCCAGTTGCATGGGCCTCATATACAATAGGATGGGAGCTGTAACCACTGAAGTGGCATTTGGCCTGGTGTGTGCAACATGTGAGCAGATTGCTGATTCCCAGCACAGGTCTCATAGGCAGATGGTGGCAACAACCAATCCATTAATAAAACATGAGAACAGAATGGTCTTGGCCAGCACTACAGCTAAGGCTTTGGTTATCCGTGTCAGGGAGCAAAAGCAGGTAGCGGAGGCCATGGAGATTGCTAGTCAGGCCAGGCAGATGGTGCAGGCAATGAGAGCCATTGGGACTTATCCGAGTCAGGGAGCAAAAGCAAGAGATGATTTTAAAGAAAAGATGCAGTCCTATAAGAAATGAACGGGGGTGCAGTTGCAACGATTCAAAAAGCCCGGATGTTGTTGCCGGGAAATTCAATGGGATCTTGCACTCAGGCTTTTAGGGACAAACTCGAACCTTGTCCCATTGATCTATGGACTCTTCAAACACGGCCGAGGAAGACGCCCTTAATGAGACGGGGGTCTTCTCAATGGGTGGAAGTTCAACTCATTCAACACCCAACATTTAAAATGTCAAGCACAAAATCATGTAGCAGACGATCAGTTTCT

Similar to Influenza A virus (A/Kenya/027/2017(H3N2)) segment 7 matrix protein 2 (M2) and matrix protein 1 (M1) genes, complete cds

### >7 H3N2 2016 Isolates from Baboons PAN 3201

TGTATACATGTTATCTTTCGCATCGTCATCAGACTCTCATGCCGAGATCGAGCAGAGACTTGCAGATGTACTTTGCTTGGTATAACACAGATCCTGAGGCTCTCATGATGACTGAGACAAGACCAATCTGTCACCTTTGACTAAGGGATTTAGGGTTTGTTTTCACGCTCACCGTGCCCAGTGAGCGAGGAGTGCAGCGTAGACGCTTTGTCCAAAATGCCCTCAATGGGAATGGAGACCCAAATAACATGGACAAAGCAGTTAAACTGTATAGGAAACTTAAGAGGGAGATAACGTTCCACGGGGCCAAAGAAATAGCTCTTAGTTATTCTGCTGGTGCACTTGCCAGTTGCATGGGCCTCATATACAATAGGATGGGAGCTGTAACCACTGAAGTGGCATTTG

GCCTGGTGTGTGCAACATGTGAGCAGATTGCTGATTCCCAGCACAGGTCTCATAGGCAGATGGTGGCAACAACCAATCCATTAATAAAACATGAGAACAGAATGGTCTTGGCCAGCACTACAGCTAAGGCTATGGTTCTAAGTGTCAGGGAACAAAAGCAGGTAGCGGAGGCCATGGAGATTGCTAGTCAGGCCAGGCAGATGGTGCAGGCAATGAGAGCCATTGGGACTTATCCGAGTTCCGGAGCAAAAGCAAGAGATGATTTTAAAGAAAATATGCAGACCTATCAGAAATGAACGGGGGTGCAGATGCAACGATTCAAGTGACCCGGATGTTGTTGCCGGGAAAATCAATGGGATCTTGCACTCGAGCTTGTGGATTCAAAATCGTCTCCTTGTCCAATGGATCTATGGACTCTTCAAACACGGCCGAGGAGACGCCCTTAATGGGCCGGAGGACCTCTCAAAGGAGGGAAGAATAACTCAAGCAACACCCAACATTTAAAATGTCAAGCCCAAAATCATGTAGCAGACGTCCTGTACCT

Similar to Influenza A virus (A/Florida/02/2018(H3N2)) segment 7 matrix protein 2 (M2) and matrix protein 1 (M1) genes, complete cds

### >8 (H1N1) 2016 Isolates from Baboons PAN3638

ATGCAAGGGTTGCGAAAACACACATGATCTCTTTGTTATACTAACACATT

ATAGACATAGTAAATTCTGTTATTGAAAAGATGAATACACAGTTCACAGC

AGTGGGTAAAGAGTTCAACCACCTTGAAAAAAGAATAGAGAATCTAAATA

AAAAAGTTGATGATGGTTTCCTGGACATTTGGACTTACAATGCCGAACTG

TTGGTTCTACTGGAAAATGAAAGAACTTTGGACTATCACGATTCAAATGT

GAAGAACTTGTATGAAAAAGTAAGAAACCAGTTAAAAAACAATGCCAAGG

AAATTGGAAACGGCTGCTTTGAATTTTACCACAAATGCGATAACACATGC

ATGGAAAGTGTCAAGAATGGGACTTATGACTACCCAAAATACTCAGAGGA

AGCAAAATTAAACAGAGAAAAAATAGATGGAGTAAAGCTGGAATCAACAA

GGATCTACCAGATTTTGGCGATCTATTCAACTGTCGCCAGTTCATTGGTA

CTGGTAGTCTCCCTGGGGGCAATCAGCTTCTGGATGTGCTCTAATGGGTC

TCTACAGTGTAGAATATGTATTTAACATTAGGATTTCAGAATCATGAGAA

AAACACCCTTGTTTCTACTAATACGAGACAGATAATAGATAATAA

Similar to Influenza A virus (A/Vacaria/LACENRS-1312/2016(H1N1)) segment 4 hemagglutinin (HA) gene, complete cds

**Appendix 1:**
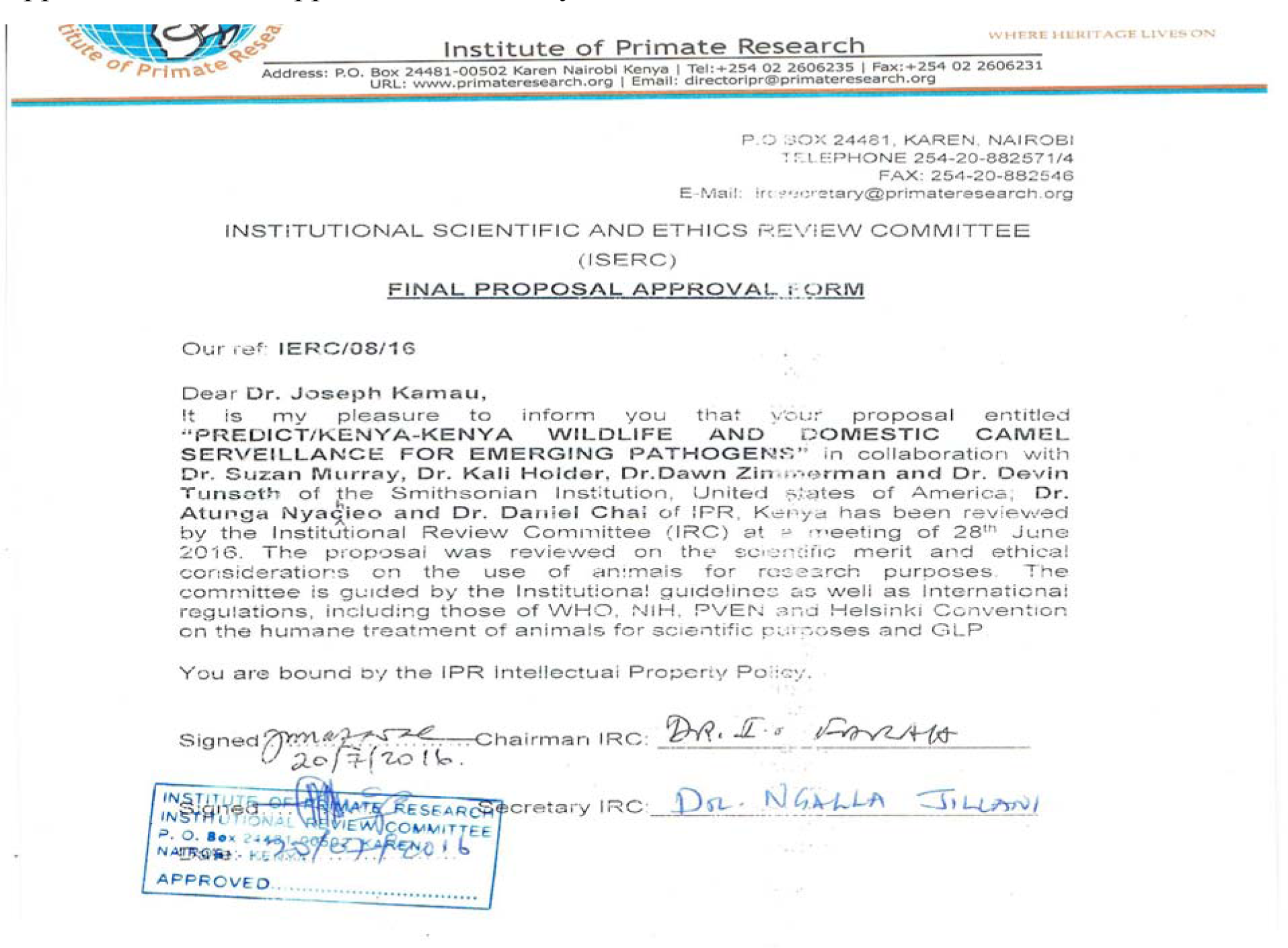
Ethical approval for the study.

**Appendix 2:**
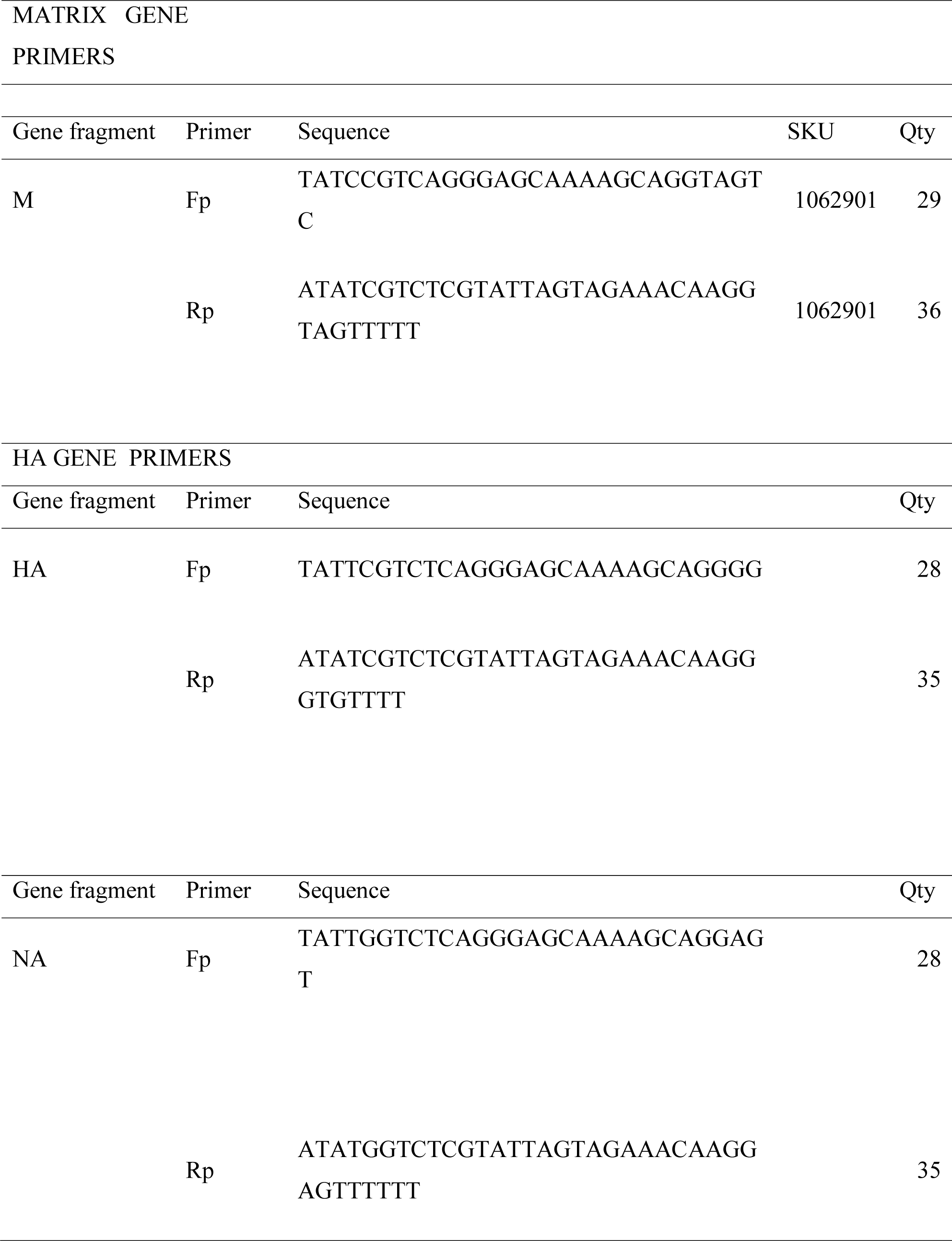
Primers used in this study

**Appendix 3.**
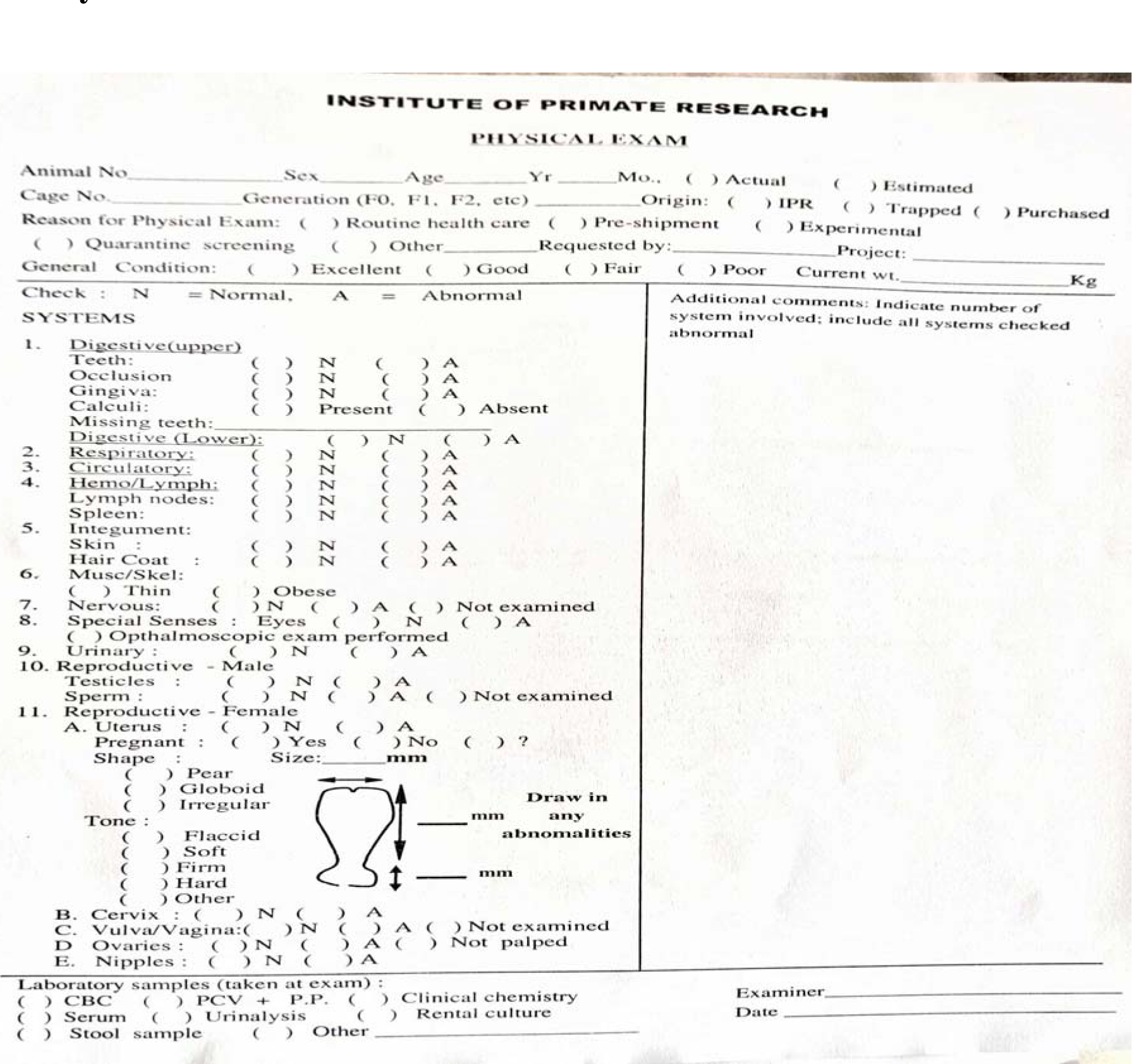
Physical examination forms

**Appendix 4:**
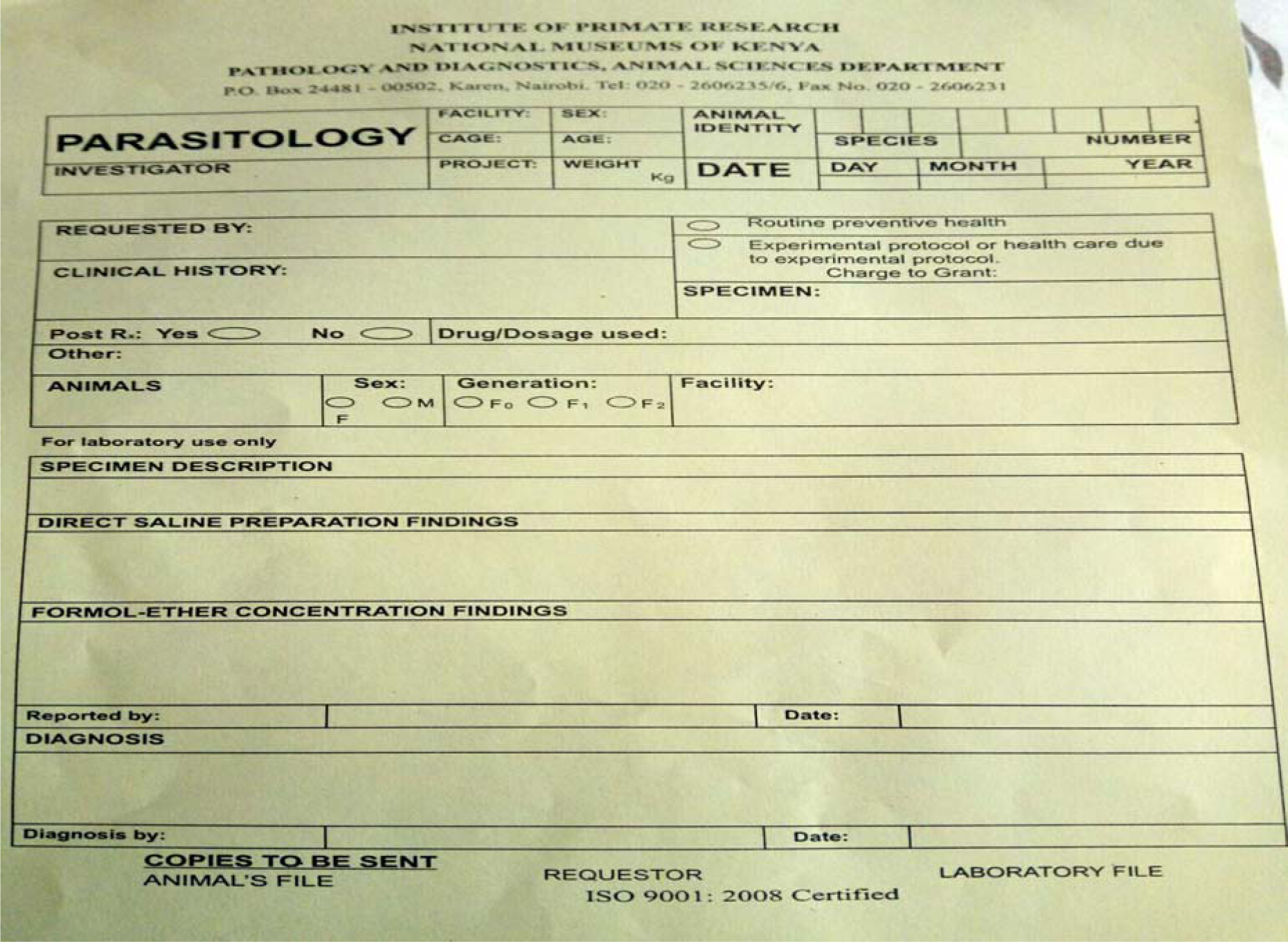
Parasitology information form

**Appendix 5:**
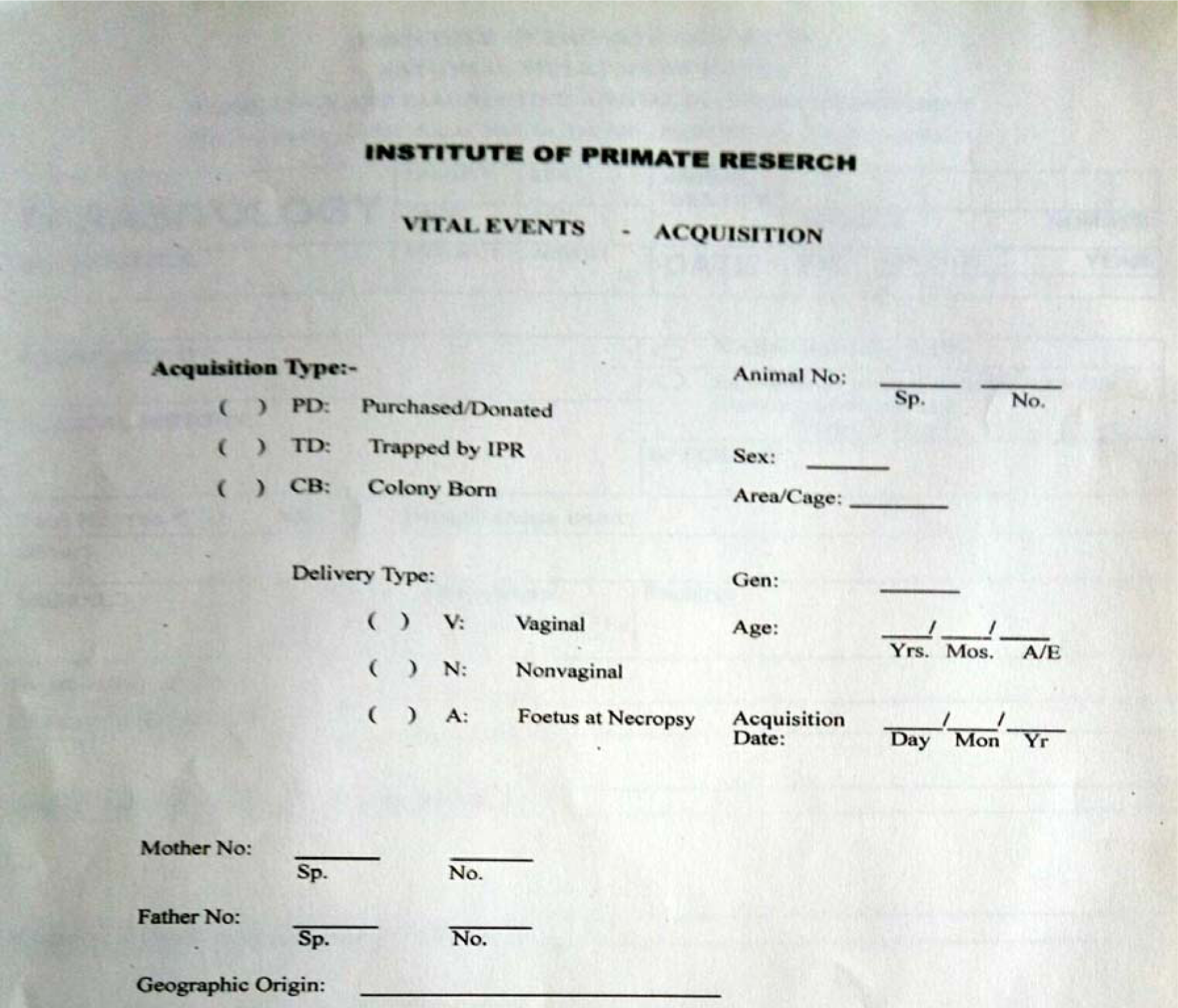
Acquisition of the animal forms for vital events

